# mRNA Localization and Local Translation of the Microtubule Severing Enzyme, Fidgetin-Like 2, in Polarization, Migration and Outgrowth

**DOI:** 10.1101/2023.04.17.537087

**Authors:** Rayna Birnbaum, Jeetayu Biswas, Robert H. Singer, David J. Sharp

## Abstract

Cell motility requires strict spatiotemporal control of protein expression. During cell migration, mRNA localization and local translation in subcellular areas like the leading edge and protrusions are particularly advantageous for regulating the reorganization of the cytoskeleton. Fidgetin-Like 2 (FL2), a microtubule severing enzyme (MSE) that restricts migration and outgrowth, localizes to the leading edge of protrusions where it severs dynamic microtubules. FL2 is primarily expressed during development but in adulthood, is spatially upregulated at the leading edge minutes after injury. Here, we show mRNA localization and local translation in protrusions of polarized cells are responsible for FL2 leading edge expression after injury. The data suggests that the RNA binding protein IMP1 is involved in the translational regulation and stabilization of FL2 mRNA, in competition with the miRNA let-7. These data exemplify the role of local translation in microtubule network reorganization during migration and elucidate an unexplored MSE protein localization mechanism.

**Graphical Abstract:** 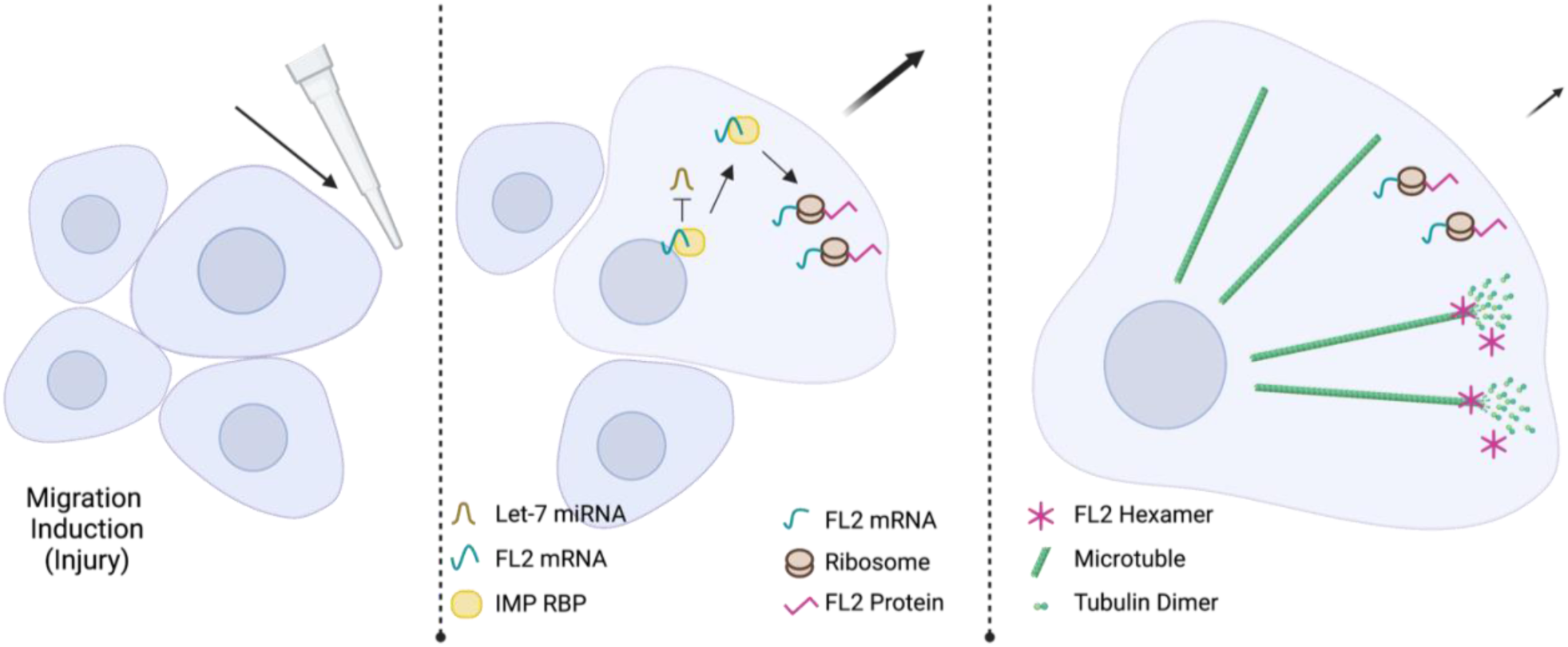

**Highlights:**

1. The microtubule severing enzyme FL2 RNA is localized to the leading edge
2. FL2 mRNA localization leads to FL2 translation within protrusions
3. The IMP family works in concert with Let-7 miRNA to regulate FL2 mRNA

## INTRODUCTION

Cellular development, polarization and motility are a few of the processes requiring specifically timed and localized protein expression. To achieve this stringent spatiotemporal control, cells utilize RNA localization and local protein synthesis. Transport of one mRNA molecule can produce many proteins in its target compartment compared to the costly process of mass transport of individual proteins, making localization of mRNA and local translation more biologically efficient, both temporally and energetically. Efficiency is only one of the many benefits of local protein synthesis, which also include rapid response to local stimuli, avoidance of ectopic interactions and controlled expression of toxic proteins^1, 2^.

Cell migration and neurite outgrowth involve the rapid reorganization of the cytoskeleton, which requires asymmetric protein expression^3^. Cell polarization and protrusion formation, the initial steps of migration, are driven by the differential buildup and breakdown of the cytoskeleton^4^. This asymmetric cellular outgrowth requires localized, asymmetric expression of cytoskeletal related proteins. The prototypical example of locally translated cytoskeletal asymmetric protein expression is β-actin, the mRNA of which was described to have strong localization to protrusions^5^, axons and growth cones^6^ during development and migration.

Actin is often thought of as the main player in cell migration and process formation, but the necessity of an intact and dynamic microtubule network to these processes is well established^7, 8^. Many studies have shown the necessity of a microtubule network for localization of mRNAs^9, 10^, but these studies focused mainly on the contribution of microtubule protein structures through association with microtubule motor proteins^11^. While emphasis has been put on regulation of actin related mRNAs^3, 12^, work on the contribution of mRNA localization and local translation to the dynamic nature of the microtubule network is still limited.

Polarization and outgrowth involve reorganization of the microtubule network. Microtubule asymmetry during motility facilitates directional polarity, intracellular transport, focal adhesion turnover, protrusion stability and growth cone turning^13–15^. While this asymmetry is well established, the involvement of local protein synthesis in the buildup and breakdown of the microtubule network has not been extensively studied.

Preliminary studies indicate that the microtubule network is regulated by local translation, similar to the actin network. The mRNA of tubulin, the main subunit of microtubules, has been identified in the peripheral cytoplasm and in axons during development and there is evidence suggesting its local synthesis in developing axons^5, 16, 17^. Microtubule associated protein (MAP1 & MAP2) mRNAs, regulators of microtubule lattice stability, have been identified in developing and adult dendrites^18, 19^. The mRNAs of the microtubule motor proteins Kinesin and Dynein were found to localize to protrusions, but evidence of local synthesis is lacking^20^.

In this study, we investigate the mRNA localization and local translation of the microtubule severing enzyme (MSE) Fidgetin Like 2 (FL2). MSEs are a class of AAA ATPase proteins involved in the process of migration^21, 22^. They use ATP hydrolysis to displace tubulin dimers within a microtubule lattice leading to depolymerization and subsequent disassembly or growth, depending on lattice stability. Regulation of microtubule dynamics via MSEs is important at the leading-edge during migration for such processes as regulation of focal adhesion assembly, microtubule lattice growth and the release of signaling molecules that affect the actin network^15^.

The Fidgetin family of MSE’s was first identified in a mutant mouse model with developmental defects in the nervous system, auditory system and skeletal structure causing “fidgeting” like symptoms^23–25^. Expression of FL2 is localized to the leading edge of migrating cells and to the growth cones of outgrowing neurites^26^. Within these subcellular areas, FL2 suppresses directed migration and outgrowth by severing the dynamic microtubules at the cell cortex and within protrusions.

FL2 is primarily expressed during development ^27, 28^, but in adulthood, FL2 is only expressed upon injury for the regulation of wound healing and regeneration. Knockdown of FL2 increases velocity of migration more than 2-fold^29^. In neurons, FL2 depletion results in increased neurite outgrowth and an attenuated response to inhibitory cues during growth cone guidance^26^. Targeting of FL2 via siRNA mediated knockdown has shown promising results for multiple therapies, including enhanced excision and burn wound healing, nerve regeneration and regeneration after corneal chemical burn^26, 29–31^.

Rapid and localized FL2 protein expression suggests that FL2 may be post-transcriptionally regulated through mRNA localization and local protein synthesis. Messenger RNA localization begins when an RNA binding protein (RBP) binds the mRNA molecule^32^. By associating with other proteins and RNAs to form a ribonucleoprotein (RNP) complex, the mRNA molecule can be either translationally repressed or activated, allowing for strict regulation of protein expression^33^. The mRNP complex can be transported to a cellular destination by binding to molecular motor proteins^34^. Once at its destination, cytoskeletal elements can anchor the complex and molecular signals, such as phosphorylation of the mRNP, can induce de-repression of the mRNA to facilitate local translation^35, 36^.

Zipcode Binding Protein 1 (ZBP1), the chicken homolog to IMP1 (Insulin like Growth Factor 2 RNA Binding Protein 1 or IGF2BP1) was the first RBP implicated in localization of β-actin mRNA^37^, the prototypical example of RNA localization to protrusions and extensions. We report here that FL2 mRNA contains the zipcode sequence^38, 39^ necessary for binding to the IMP RBPs. Furthermore, FL2 was previously identified as an mRNA target of let-7 microRNA^40^, short, non-coding RNAs that decrease expression of their target, either by mRNA degradation or translational suppression. Recent evidence has shown that IMP RBPs and let-7 miRNAs have an antagonistic relationship; the IMP RBPs can protect mRNAs from let-7 mediated downregulation, and the IMP RBP mRNAs are themselves a target of let-7 miRNAs^41^. These data suggested that FL2 expression may be regulated by an antagonistic relationship between the IMP RBPs and let- 7 miRNAs.

In this work, we elucidate the post-transcriptional regulation of the MSE FL2. Using single molecule imaging, we found that FL2 mRNA is localized to and translated at the leading edge of polarized cells. This data suggests mRNA localization and stabilization by IMP1 RBP, and protection from Let-7 mediated miRNA degradation. This study provides insight into the mechanism of FL2 expression and elucidates a unique mechanism for regulating MSE localization. These data advance our understanding of cytoskeletal regulation in migration and the cellular response to injury.

## RESULTS

### FL2 protein was upregulated after injury

FL2 protein expression is minimal in adulthood but is significantly upregulated upon injury^26, 29–31^. FL2 protein is localized to the leading edge and cell cortex where FL2 carries out its severing function on dynamic microtubules during migration and outgrowth^29^. However, the specific time course of FL2 upregulation on a cellular level had yet to be determined.

To investigate the time scale of leading edge FL2 protein expression, we utilized an in vitro wound healing scratch assay to induce directional migration. Compared to cells in an uninjured monolayer, FL2 protein fluorescence intensity at the leading edge increased significantly (p=0.0005) in cells on the wound periphery just five minutes after injury (Fig 1A, F). FL2 protein concentration in the leading edge continued to increase for one hour after injury (Fig 1B, F). These data indicated that FL2 protein is either made or rapidly reorganized as soon as five minutes after injury and protein concentration in this compartment continues to increase over time in response to migration induction.

**Figure 1.**
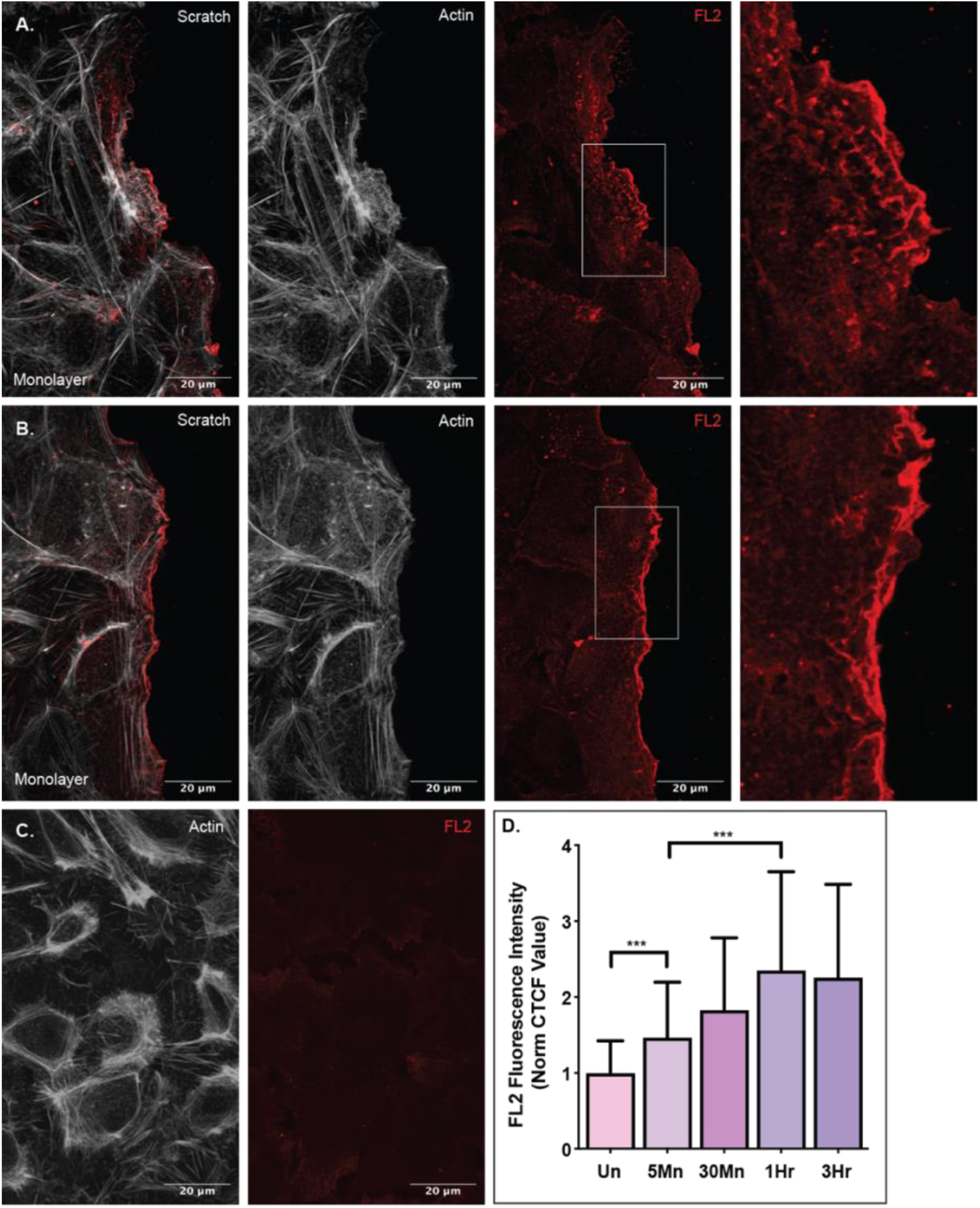
FL2 expression at the leading edge increases over time upon migration. FL2 protein stain (red) and actin network stain (white) in U2OS cells incubated after an in vitro scratch injury. A) Expression of FL2 protein and the actin network in cells fixed five minutes after a scratch injury. Sequence of images is a merged image of FL2 and actin, an image of the actin stain, an image of the FL2 stain and a zoomed in image of the cell edge indicated by white box in previous image. B) Expression of FL2 protein and the actin network in cells fixed on hour after injury. Sequence of images is the same as panel A. C) Expression of FL2 protein and the actin network in an uninjured cell monolayer. D) Quantified FL2 fluorescence intensity of the leading edge (first five micrometers of the cell facing the scratch zone) of cells on the scratch zone periphery of cells incubated for 5Mn (five minutes; n=42), 30Mn (thirty minutes; n=40), 1Hr (one hour; n=41) and 3Hr (three hours; n=40) or the quantified fluorescence intensity of the cortex of cells in an Un (uninjured monolayer; n=41).

### FL2 protein at the leading edge was newly synthesized upon injury

Local protein synthesis has many advantages, including rapid induction of highly localized protein expression without the energy cost and time of transporting the protein. Increase in localized FL2 protein expression just five minutes after induction suggested that FL2 protein may be locally translated.

To determine whether FL2 translation occurred upon injury, we performed in vitro scratch assays on cells pre-treated with the translation inhibitor puromycin. As indicated in Figure 1, one hour after injury was the maximum of short term FL2 protein expression on the cellular level. Thus, the monolayers were incubated for one hour after injury with and without puromycin treatment. The number of cells on the wound periphery that had significant FL2 upregulation at the leading edge were quantified. Significant FL2 upregulation was defined as FL2 fluorescence intensity that was 40% greater within the leading edge than in the remaining cytoplasm. Compared to untreated coverslips (Fig 2A), there was a 50% decrease in number of cells with significant FL2 protein upregulation at the leading edge on coverslips treated with puromycin (Fig 2B & 2C). This significant decrease (p<0.0001) in the number of cells with injury induced FL2 protein upregulation at the leading edge indicated that much of the protein expression was newly synthesized upon polarization and migration induction, further adding to the evidence of local FL2 translation.

**Figure 2.**
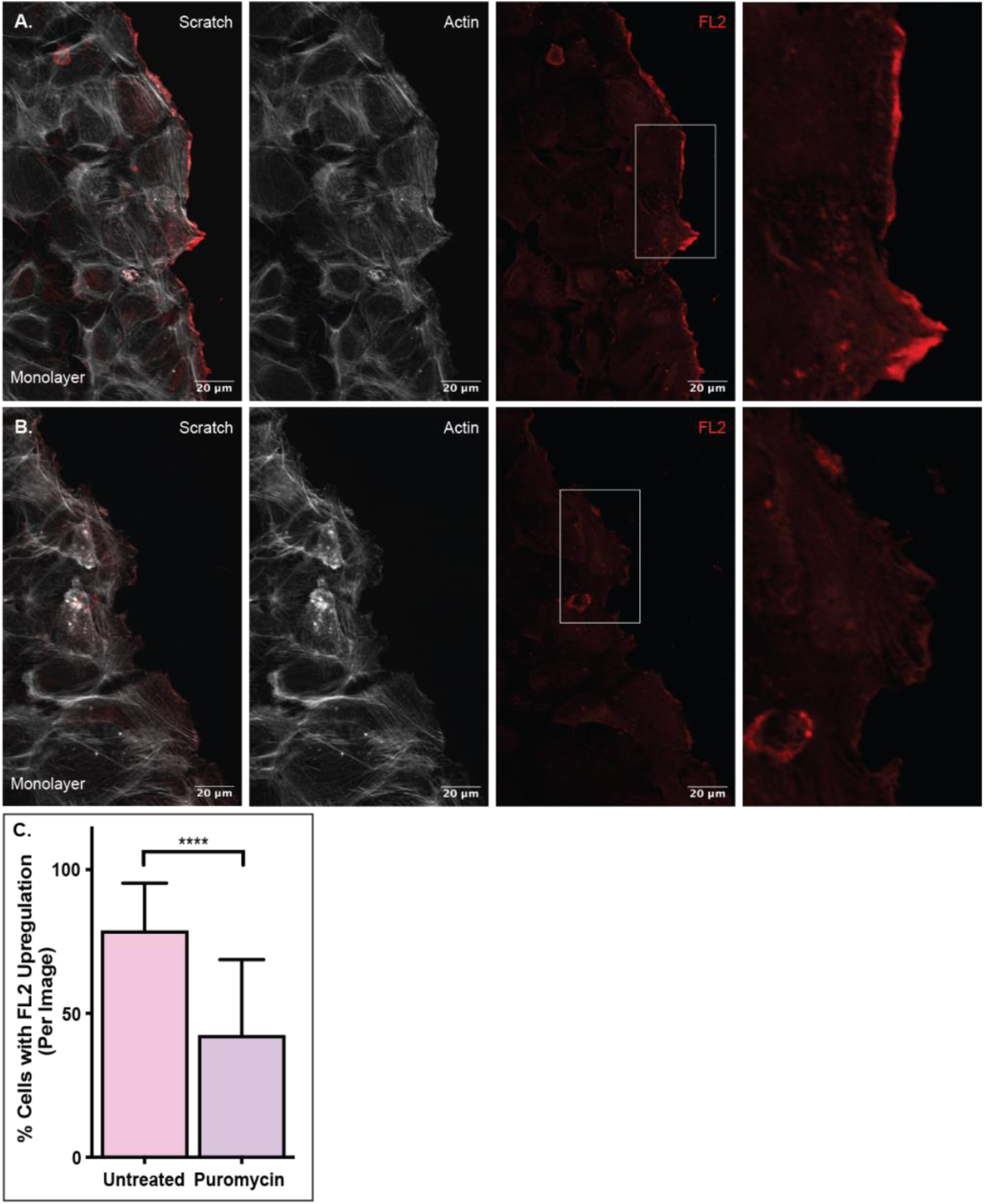
FL2 protein is newly synthesized upon migration. FL2 protein stain (red) and actin network stain (white) in U2OS cells incubated for one hour after a scratch. A) FL2 protein expression in untreated cells after a scratch. Sequence of images is a merged image of FL2 and actin, an image of the actin stain, an image of the FL2 stain and a zoomed in image of the cell edge indicated by white box in previous image. B) FL2 protein expression in cells treated with 200ug of puromycin prior to and during scratch. Sequence of images is the same as panel A. C) Quantification of the number of cells with significant FL2 protein expression at the leading edge. Untreated n=63; Puromycin n=63. Significant protein expression defined as fluorescence intensity 40% greater than the cytoplasm of the same cell and spanning at least a fourth of the cell edge.

### FL2 mRNA was localized to the leading edge and growth cones

For local translation to occur in a specific cellular compartment, the mRNA must first localize there. To investigate if FL2 mRNA was localized to the leading edge, we utilized a single molecule fluorescent *in situ* hybridization (smFISH) assay that was optimized for the high GC content (74%) of FL2 mRNA. This method used an initial primary set of probes to hybridize to the sequence of interest and a secondary, fluorescently labeled set of probes to hybridize to the primary probes^42^. The optimized approach led to higher fidelity of detection and decreased background when compared to traditional smFISH (Sup Fig 1).

In U2OS cells one hour after scratch injury, an average of 45% of the FL2 mRNA molecules localize to the leading edge of protrusions in polarized cells on the wound periphery (Fig 3C & D). The result is particularly noteworthy when considering the area (Fig 3A) and volume of the leading edge, a thin membrane protrusion, compared to that of the remaining cytoplasm.

**Figure 3.**
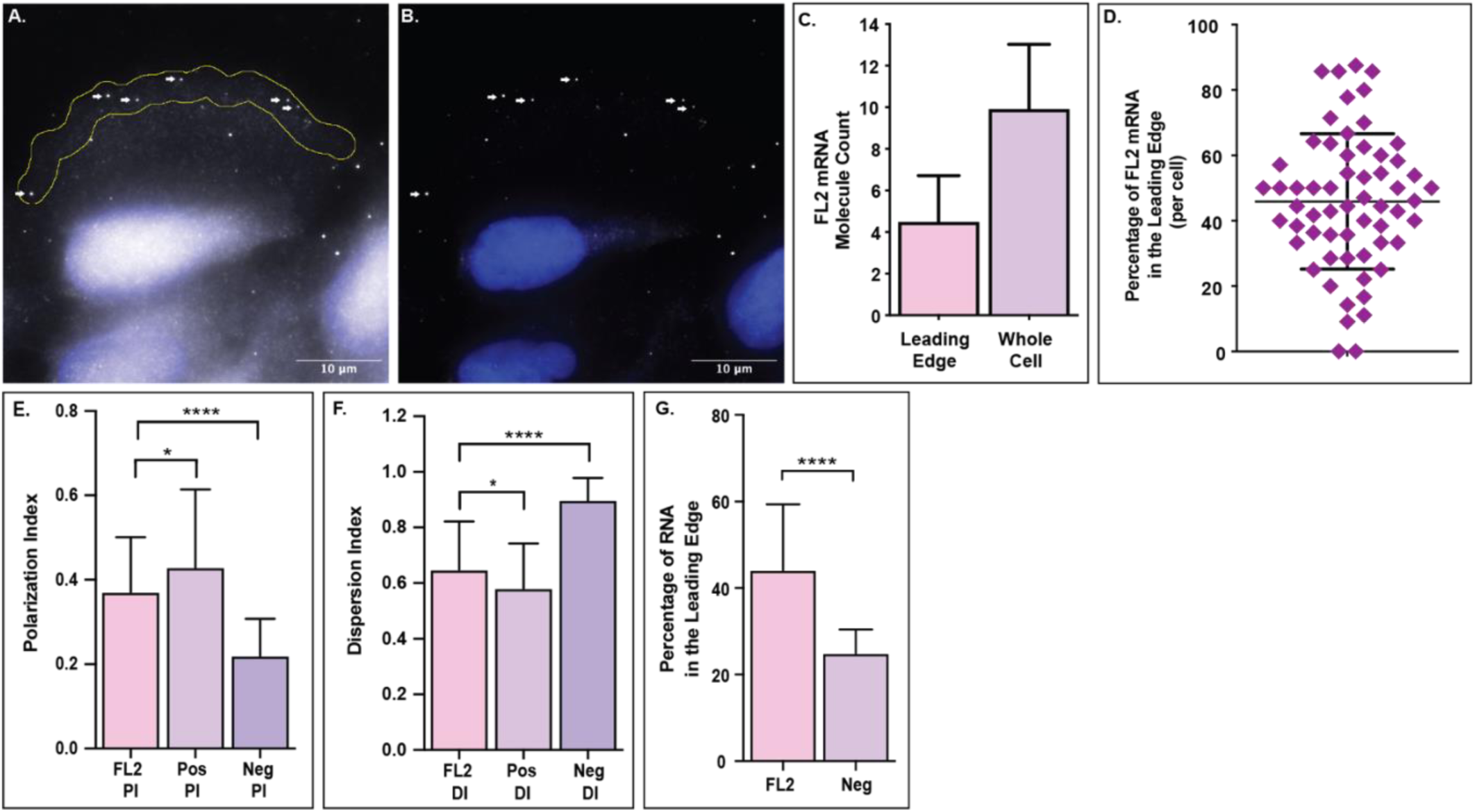
FL2 mRNA localizes to the leading edge of polarized cells. FL2 mRNA in U2OS cells one hour after a scratch injury. A) Image of FL2 mRNA molecules (white) in a polarized cell on the scratch periphery. Yellow region indicates the leading edge, defined as the first five micrometers of the cell facing the scratch zone. White arrows indicate the FL2 mRNA molecules within the leading edge. DAPI nuclear stain in blue. B) Filtered image of the same cell from panel A to decrease background and emphasize FL2 mRNA localization within the cell. White arrows indicate the same FL2 mRNA molecules as in panel A. C) Quantification of FL2 mRNA within the leading edge (indicated by the yellow outline in panel A) compared to the amount of FL2 mRNA within the whole cell (n=60). D) Percentage of FL2 mRNA within the leading edge of individual cells on the scratch periphery. E) Polarization index (PI) of FL2 mRNA (n=98 pooled) compared to the PI of a positive control (Pos; β-actin mRNA; n=65) and negative control (Neg; IMP2 mRNA; n=33). F) Dispersion index (DI) of FL2 mRNA compared to the DI of a positive control (Pos; β-actin mRNA) and negative control (Neg; IMP2 mRNA). Same cells at panel E. G) Leading edge analysis of FL2 mRNA compared to a negative control (Neg; IMP2 mRNA) in the same cells. N=31.

As a secondary mRNA localization analysis, we compared FL2 mRNA localization to the localization of positive (β-actin mRNA; Pos) and negative (IMP2 mRNA; Neg) controls using the previously published polarization and dispersion index assay^43^ (Sup Fig 2). FL2 mRNA was less polarized (p=0.0197) than the positive control, but significantly more polarized (p<0.0001) than the negative control (Fig 3E). FL2 mRNA was more disperse (p=0.0148) than the positive control, but significantly less disperse (p<0.0001) than the negative control (Fig 3F). We then compared the leading edge analysis of FL2 mRNA to that of the negative control. FL2 mRNA localized to the leading edge (45%) at a rate significantly higher (p<0.0001) than the negative control mRNA (25%) (Fig 3G). Taken together, these data showed that FL2 mRNA was asymmetrically distributed with preferential localization to the leading edge of polarized cells.

Similar localization of FL2 mRNA was seen in the protrusions of NIH3T3 mouse embryonic fibroblasts, protrusions of human BeWo cells and in the extensions and growth cones of differentiating mouse neuro2A cells and human iPS derived neuronal cells (Fig 4). These data support the notion that FL2 mRNA localized to the leading edge of protrusions and extensions, the same cellular compartments where localized FL2 protein expression occurs.

**Figure 4.**
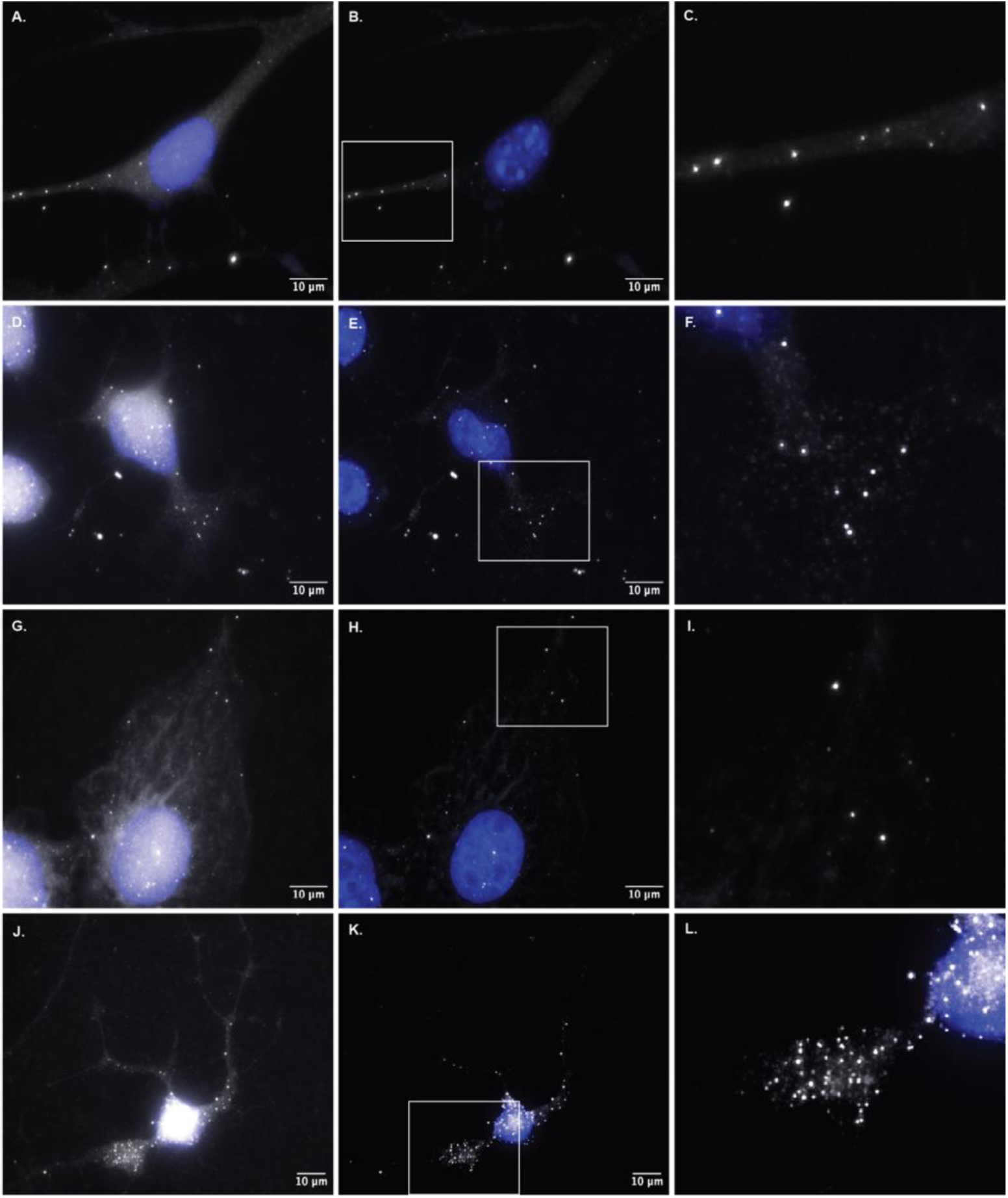
Localization of FL2 mRNA to protrusions and growth cones of different cell types. Optimized single molecule fluorescence *in situ* hybridization in different cell types for FL2 mRNA. A) FL2 mRNA in a polarized NIH3T3 mouse embryonic fibroblast (MEF). B) Filtered image of the cell in panel A. C) Zoomed in image of FL2 mRNA localized to the extension of the MEF from panels A & B. Area indicated by the white box in panel B. D) FL2 mRNA in a mouse Neuro2A (N2A) neuroblastoma cell differentiated for 48 hours prior to fixation. E) Filtered image of the N2A cell in panel D. F) Zoomed in image of FL2 mRNA localized to the growth cone of the N2A cell from panels D & E. Area indicated by the white box in panel E. G) FL2 mRNA in a human BeWo choriocarcinoma placenta cell. H) Filtered image of the BeWo cell in panel G. I) Zoomed in image of FL2 mRNA localized to the edge protrusion of the BeWo cell in panels G & H. Area indicated by the white box in panel H. J) FL2 mRNA in a human induced pluripotent stem (iPS) derived neuronal cell differentiated for five days prior to fixation. K) Filtered image of the neuronal cell from panel J. L) Zoomed in image of FL2 mRNA localized to the growth cone of the neuronal cell from panels J & K. Area indicated by the white box in panel K.

### Imaging of FL2 translation in cellular protrusions using the SunTag

The translation of FL2 protein after injury and the presence of FL2 mRNA in the leading edge and in neuronal processes suggested that FL2 was being newly translated in these cellular compartments. To directly visualize local FL2 translation, the SunTag system was utilized to perform single molecule imaging of nascent peptides^44^.

Colocalization of the 24x GCN4 peptide repeats of the SunTag with single mRNA molecules allowed for the visualization of translating proteins^44, 45^. FISH-IF in U2OS cells expressing a construct with the SunTag fused to the N terminus of FL2 (CDS and 3’UTR; Fig 5A) identified individual translation sites. SunTag (GCN4) immunofluorescent stains were imaged along with the optimized smFISH of FL2 mRNA probes. Colocalization of the SunTag stain and FL2 mRNA probes was determined, indicating sites of translation (Fig 5B – J). FL2 mRNA spots and translation sites were binned based on location; either internally around the nucleus (5um out from the DAPI nuclear stain) or more peripherally (further away from the nucleus, including cellular protrusions). Quantification of both the FL2 mRNA spots and translation sites showed that there was a higher percentage of translating mRNA in the peripheral region (average of 31%) compared to the internal region surrounding the nucleus (average of 12%) in each cell (Fig 5K). An FL2 mRNA molecule was about 3-fold more likely on average to be translating if it was localized to the peripheral region compared to the perinuclear region of the cell (Fig 5L). These data indicated that the translation of FL2 mRNA occurred within protrusions of polarized cells.

**Figure 5.**
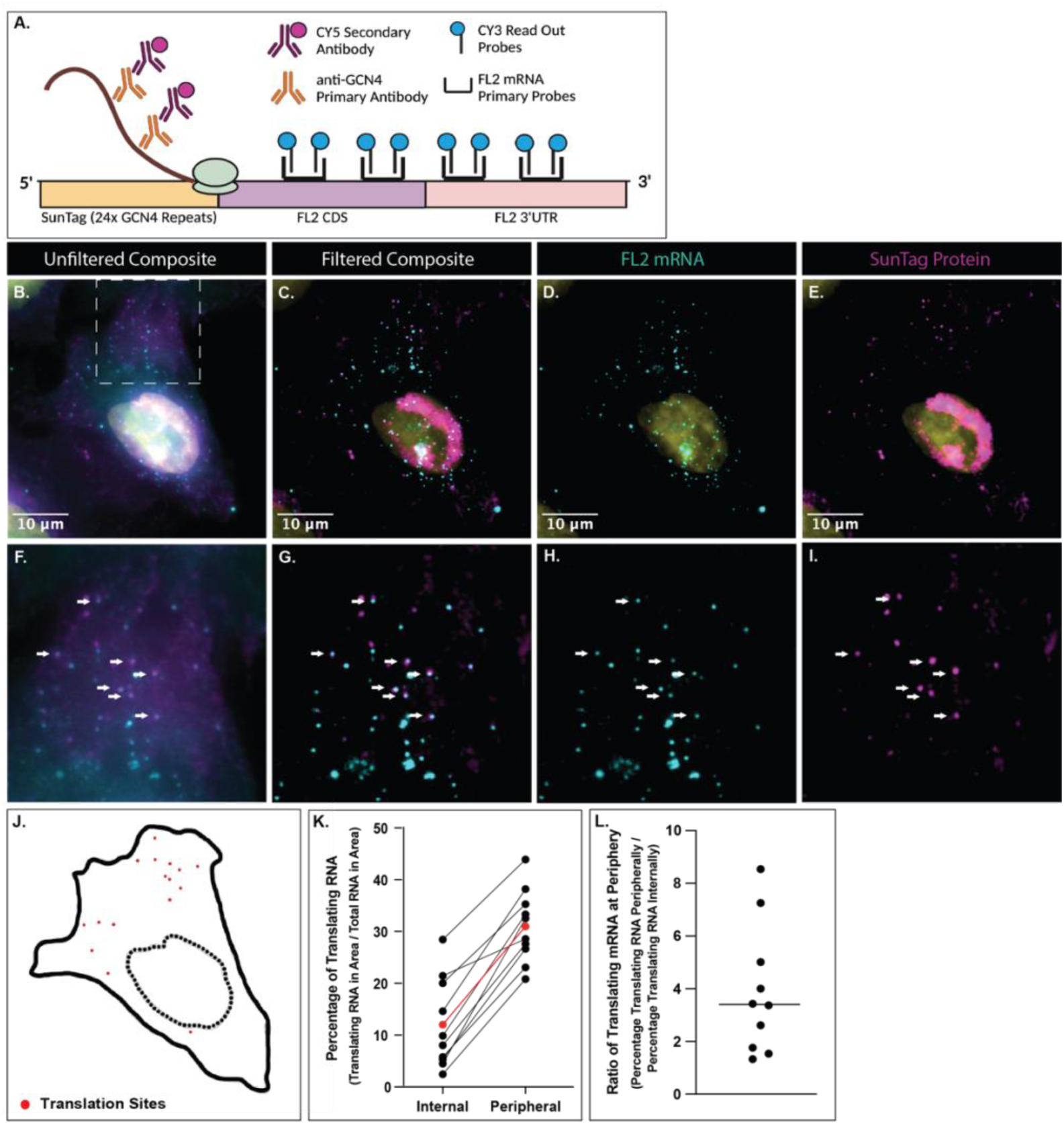
FL2 is locally synthesized within the periphery of polarized cells. U2OS cells expressing a Ubc:SunTag:FL2:FL2(3’UTR) construct. A) Schematic of the SunTag:FL2:FL2(3’UTR) construct used to image translation of FL2. The anti-GCN4 primary antibody binds to the translating GCN4 protein repeats while optimized smFISH probes bind to the FL2 mRNA sequence. Colocalization of the CY5 secondary antibody and the CY3 smFISH probes indicates a site of translation. B) Composite of the CY5 and CY3 unfiltered images of a U2OS cell expressing the SunTag:FL2:FL2(3’UTR) construct. White box indicates the region shown in panels F - I. C) Composite of the CY5 and CY3 filtered images of the cell shown in panel B. D) Cy3 (FL2 mRNA) filtered image of the cell in panel B & C. E) CY5 (SunTag protein) filtered image of the cell in panel B & C. F - I) Zoomed in images of panels B - E respectively. Area indicated by white box in panel B. White arrows pointing to sites of colocalization determined via FISHQUANT^72^, indicating translation sites. J) Outline of the cell in above panels. Dotted line indicating outline of DAPI stain (nucleus). Red dots in the location of colocalization of CY3 (FL2 mRNA) and CY5 (SunTag protein) which indicates translation sites. K) Percentage of translating RNA in the internal region and peripheral region of individual cells. Each black line is a single cell (n = 10). Red line is the average percentages of all cells. L) The ratio of translating mRNA at the periphery compared to those internally. The line indicates that the average peripheral mRNA is three times more likely to be translating.

### Post-transcriptional regulators implicated in FL2 mRNA expression

As localization of FL2 mRNA at the leading edge was similar to that of β-actin mRNA^46^, the prototypical example of leading-edge mRNA localization, it was possible that they may be regulated by similar mechanisms. Prior work has extensively characterized the zipcode sequences in RNA necessary for binding to the IMP family of RBPs^36, 39^. We identified the presence of zipcode like sequences in both human and mouse FL2 mRNA sequences (Fig 6A).

**Figure 6.**
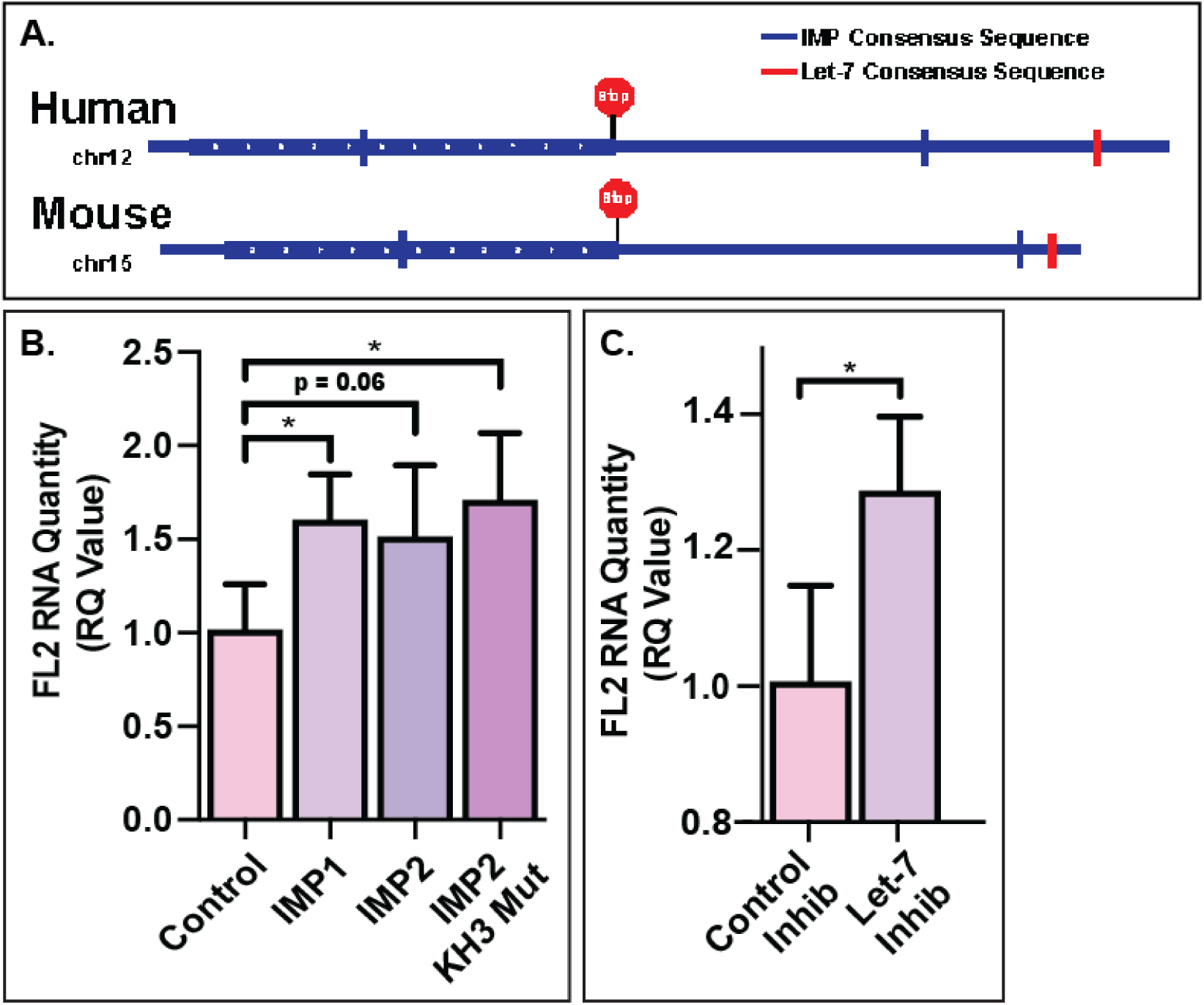
IMP RBPs and let-7 miRNA are involved in the regulation of FL2 mRNA. A) Representation of the IMP RBP and let-7 miRNA binding sites on the FL2 mRNA sequence. B) FL2 mRNA in N2A cells with 48 hour IMP RBP overexpression quantified via RT-PCR. C) FL2 mRNA in N2A cells treated with 48 hour let-7 miRNA inhibition.

As IMP RBPs have been shown to affect localization^37, 47^, translation^32, 48^ and degradation^49^ of their targets, RT-PCR was used to quantify FL2 mRNA levels in N2A cells transfected with IMP RBP (IMP1, IGF2BP1 and IMP2, IGF2BP2) plasmids. Overexpression of both IMP1 (p=0.0134) and IMP2 (p=0.0672) increased the FL2 mRNA levels compared to transfection control (pMax-GFP). To determine whether FL2 had a higher binding affinity to IMP1 compared to IMP2, an IMP2 mutant was used that caused IMP2 to recognize IMP1 binding sequences^39^. Overexpression of the IMP2 mutant rescued the level of FL2 mRNA (p=0.0174), increasing it to a level comparable with that of IMP1 overexpression (Fig 6B). These data suggested that both IMP1 and IMP2 may bind FL2 mRNA and protect it from degradation, with a higher binding affinity to IMP1.

The increase in FL2 mRNA levels with IMP overexpression suggested that the IMPs may be protecting FL2 mRNA from miRNA mediated degradation, as has been shown for other IMP targets such as HMGA2^41, 50, 51^. To investigate the role of miRNAs in the regulation of FL2 mRNA, the sequences of human and mouse FL2 were investigated for seed sequence matches to miRNAs, which identified the Let-7 family of miRNAs (Fig 6A). A let-7 miRNA family inhibitor^52^ significantly increased the level of FL2 mRNA (p=0.0192) compared to a control inhibitor (Fig 6C), similar to the observed increase with overexpression of the IMP RBPs.

A negative regulator of let-7 miRNA, LIN28 RBP, targets the let-7 precursors. Previous studies have identified the importance of the direct or indirect interactions of the let-7 miRNA, IMP RBPs and LIN28 RBP on target mRNAs^41, 49, 51^. Overexpression of LIN28A in N2A cells increased (p=0.0199) levels of FL2 mRNA quantity (Sup Fig 3). To determine if this increase in FL2 mRNA was due to a direct interaction, a LIN28A mutant that does not bind to the let-7 precursor was tested. Overexpression of the LIN28A mutant showed no change in FL2 mRNA levels, indicating that LIN28A had an indirect effect on FL2 mRNA through the downregulation of let-7 miRNA.

Increased quantity of FL2 mRNA with let-7 inhibition indicated that the let-7 miRNA family was likely involved in the regulation and degradation of FL2 mRNA expression. Along with the increase in FL2 mRNA seen with overexpression of the IMP1 RBP, these data suggested that FL2 mRNA was locally translated like other IMP1 targets and protected by IMP1 from let-7 miRNA mediated degradation.

## DISCUSSION

Using translation inhibition and single molecule imaging, we confirmed that FL2 mRNA was locally translated at the leading edge of protrusions during polarization, migration and outgrowth. This data suggested this process was regulated by both IMP RBPs and the miRNA let-7, which play antagonistic roles.

### Local translation in cytoskeletal reorganization

Migration and outgrowth are cellular processes that require strict temporal and spatial regulation of protein expression, particularly important for cytoskeletal reorganization. It is well known that microtubule proteins are important in motility; cellular motility cannot occur without an intact and dynamic microtubule array^13^. However, little work has focused on the local translation of proteins involved in the microtubule network.

Many studies investigating cytoskeletal associated mRNA localization and subsequent local protein synthesis have revolved around actin network related proteins, such as β-actin, arp2/3, cofilin 1 and small Rho GTPases ^5, 6, 53–56^. However, evidence of localization and local translation of microtubule related mRNAs is growing ^5, 16–20^. In neurons, β-III tubulin expression in developing axons was decreased after local treatment with a translation inhibitor, suggesting local translation^16, 57^. Additionally, some mRNAs of microtubule accessory proteins, such as Tau, MAP1 and MAP2 mRNAs, have been identified in axons and dendrites^10, 18, 58^. Here we show the mRNA localization and local translation of a microtubule remodeling protein that is involved in microtubule dynamics during migration. This study provides further evidence that local translation is important in microtubule reorganization during motility.

### Regulation of microtubule severing enzymes through local translation

All microtubule severing enzymes (MSEs) have specific spatiotemporal expression patterns. Katanin, a well-studied MSE, localizes to protrusions, microtubule minus ends and is concentrated at spindle poles^59^. Katanin is also required for cytokinesis and, along with Fidgetin and Spastin, plays a role in poleward flux during mitosis, dendritic pruning and axonal growth during development^21^. These are just a few examples of the necessity of spatial and temporal regulation of MSEs for proper cellular growth and motility.

Most MSE protein localization studies focus on post-translational protein targeting through recognition of specific microtubule structures, microtubule associated proteins or tubulin post- translational modifications^22, 60, 61^. A recent study into transcriptional and translation regulation of MSE expression discovered transcriptional and post-transcriptional regulation of Katanin and Spastin by the transcription factor Elk1 and translational repression by the mRNA binding protein HuR^62^.

This study proposes a possible contribution of local translation to MSEs protein localization. Here we showed that the localization of FL2 protein to the leading edge was driven by mRNA localization and local protein synthesis in the same compartment. Additionally, aside from FL2, binding sites for IMP1 (ZBP1) and IMP2 RBPs were identified in the gene sequences of fidgetin, fidgetin-like1, katanin and spastin using the motif finder of Integrative Genomics Viewer (Sup Fig 4). These data implicate mRNA regulation and local translation in the spatiotemporal regulation of all MSEs.

### Implications in growth cone collapse

Previous studies have identified FL2 as a regulator of migration and outgrowth by severing the dynamic microtubules within the leading edge of protrusions and within growth cones. Specifically in growth cones, FL2 expression is involved in guidance and collapse in response to inhibitory cues, such as Semaphorin 3A and Chondroitin Sulfate Proteoglycans. Knockout of FL2 results in a significantly diminished response to these repulsive guidance cues^26^. A similar diminished response to repulsive guidance cues occurs with translation inhibition in isolated growth cones^63, 64^. Taken together with our data, these results raise the possibility that the regulation of FL2 in the growth cone is dependent upon local translation.

Recent findings from our lab indicated that FL2 activity in protrusions caused the release of GEFH1, which then locally activates RhoA^65^. RhoA is a small GTPase that is also locally translated in axons in response to inhibitory cues, where it stimulates the retraction of the growth cone^66^. Altogether, these data suggest that FL2 local translation in response to repulsive cues could lead to the stimulation of the simultaneously locally translated RhoA to induce growth cone collapse and retraction. Hence, this work provides evidence for a possible downstream pathway that induces growth cone collapse and directional outgrowth restriction.

### RNA binding proteins and miRNAs in FL2 regulation

The IMP family of RBPs is important in mRNA localization to the leading edge and growth cones, particularly in development^67^. IMP containing ribonucleoprotein (mRNP) granules can be found within protrusions and in growth cones during migration and neurite outgrowth^49^. Within these granules, IMP1 has been shown to bind many cytoskeletal mRNAs that are destined for protrusions and extensions, including the mRNAs of β-actin, Arp2/3, E-cadherin, and alpha actinin^56, 68^. Like IMP1, FL2 expression is primarily developmental and plays a major role in migration and outgrowth. The local translation of FL2 within protrusions is consistent with the role of the IMPs in motility, particularly during development.

This study also identified let-7 miRNA as a regulator of FL2 expression. Recently, the IMP RBP, LIN28 and let-7 miRNA triad has been investigated for its importance in development. The target mRNA transcripts are either bound by the IMP RBP, directly preventing the binding of let-7 miRNA, or sequestered into an IMP containing mRNP complex lacking let-7 miRNA and its degradation machinery^41, 49, 51^. LIN28 can indirectly effect expression of let-7 miRNA targets by preventing the maturation of the let-7 precursor and thereby decreasing its expression. Since an FL2 knockout is embryonic lethal, the increase in FL2 mRNA quantity with IMP overexpression, let-7 inhibition and LIN28 overexpression was consistent with previous studies implicating this triad in proper developmental progression.

### Limitations of the study

Limitations in immunofluorescent (IF) stains and qPCR efficiency have restricted experimentation of FL2. There is no validated IF stain for mouse FL2 protein. Additionally, multiple labs have been unsuccessful in attempts to use qPCR to quantify changes in human FL2 mRNA expression, most likely due to the high GC content of the sequence. Due to these shortcomings, multiple different cell lines were used throughout this study. The consistency of FL2 mRNA localization in multiple cell types of both human and mouse indicated that FL2 mRNA regulation mechanisms were similar across species.

This study indicated the involvement of IMP RBPs and let-7 miRNAs in FL2 mRNA regulation, but did not demonstrate direct binding. The qPCR analysis showed a change in FL2 mRNA expression with changes in the expression of the IMP RBPs and let-7 miRNA, but an assay specific for binding would be necessary to definitively determine whether these molecules directly interact. Further research will elucidate the specific interactions of the proteins and RNAs involved in the regulation of FL2 expression.

## Conclusions

Altogether, this work describes the spatiotemporal regulation of expression of a protein important in development and motility, and in wound healing and regeneration. This study determined that FL2 was being translated in the same cellular compartment as its localized protein expression.

Positive results from translational studies establish FL2 as a promising target for wound healing and regeneration therapeutics. Understanding the regulatory mechanisms of FL2 expression could enhance therapeutic development. Aside from the translational implications, knowledge of FL2 regulation also furthers understanding of wound healing and regeneration on a cellular level and the role that the microtubule network and supporting proteins play.

## ACKNOWLEDGMENTS

We thank Erika Pedrosa and the laboratory of Dr. Herb Lachman for the donation of human derived iPS cells. We would like to thank Dr. Carolina Eliscovich for supplying us with the stellaris FISH probes against FL2 and we would like to thank Dr. Sulagna Das and members of the Singer Lab for discussion and feedback regarding the SunTag imaging experiments performed within this study. We are grateful for the editorial contributions of Dr. Adam Kramer and Dr. Lisa Baker. We thank Andrea Briceno and the Albert Einstein College of Medicine Analytical Imaging Facility, partially funded by NCI Cancer Center Support Grant P30CA013330. Graphical abstract created with BioRender.com. J.B. was supported with funding from an MSTP Training Grant T32GM007288 and predoctoral fellowship F30CA214009. This work was supported by the NIH (5R01DK109314-05 and 5R42DK117684-03 to D.S. and R01NS083085 to R.H.S.).

## AUTHOR CONTRIBUTIONS

Conceptualization, R.B., J.B., D.J.S. and R.H.S.; Methodology and Investigation, R.B. and J.B.; Software, J.B.; Writing – Original Draft, R.B.; Writing – Review & Editing, R.B., J.B., D.J.S. and R.H.S.; Funding Acquisition and Supervision, D.J.S. and R.H.S.

## DECLARATION OF INTERESTS

The authors declare no competing interests.

## METHODS

### Cell culture, treatment, and transfection

#### Cell culture

U2OS, BeWo and N2A cells were purchased from ATCC (HTB-96, CCL-98, CCL-131 respectively). U2OS and N2A cells were cultured in DMEM supplemented with 10% fetal bovine serum, 1% Glutamax. BeWo cells were cultured in the ATCC-formulated F12-K Medium supplemented with 10% fetal bovine serum. All cells were cultured at 37 °C in the presence of 5% CO2.

#### Induced Pluripotent Stem Cell derived neurons

IPSC derived glutamatergic neuronal cells were donated to our laboratory by Erika Pedrosa from the laboratory of Dr. Herbert Lachman. Induction of the neuronal cells was achieved by forced Neurogenin-2 overexpression as previously described^69^. Neuronal cells were fixed after three days of differentiation.

#### In vitro scratch injury model

U2OS cells were grown to a confluent monolayer on glass coverslips in 24 well plates. The media in the well was aspirated and a 200ul pipette tip was used to physically scratch a “wound zone” through the monolayer. The coverslip was then rinsed with PBS to remove debris before adding fresh media back into the well. Prior to fixation, scratches were incubated for the indicated amount of time under normal culture conditions described above.

#### Transfections

N2A cells were seeded at 300,000 cells per well in a 6 well plate 24 hours before transfection using Lipofectamine 3000 (Thermo Fisher Scientific). Transfections were performed in accordance with the Lipofectamine 3000 protocol. The media was changed 24 hours after transfection to minimize cell death.

### Immunofluorescence

#### Fixation and staining

Cells were washed with PBS and incubated at 37 °C for 5 minutes in warm fixation solution consisting of 4% paraformaldehyde, 0.1% Triton X and 0.15% glutaraldehyde in BRB80. Cells were then rinsed with PBS before incubation in 10mg/mL sodium borohydrite in PBS for 20 minutes, followed by a PBS rinse and permeabilize with 0.5% TritonX in PBS for 5 minutes. Cells were rinsed with PBS supplemented with 0.05% Tween (PBST) prior to a one hour, room temperature incubation in blocking buffer (5% normal goat serum, 2mM sodium azide and 0.1% TritonX in PBS). After blocking buffer incubation, cells were incubated overnight at 4 °C in the primary antibody diluted in the blocking buffer. On the second day, the cells were washed in PBST for 5 minutes thrice before a 1.5 hour incubation in the secondary antibody diluted in the blocking buffer. Cells were washed thrice more for 5 minutes with PBST prior to mounting of the coverslips.

#### Antibodies

Monoclonal FL2 (FIGNL2; NM_001013690) antibody generated in mice (Abmart, Shanghai, China) using an antiFL2 (SSTTPSPAHK).

#### Alexa Fluor 488 Phalloidin (Invitrogen #A12379)

Rabbit anti-GCN4 (SunTag) antibody (absolute antibody #Ab00436-230) anti-Mouse Secondary Antibody, Alexa Fluor 568 (Invitrogen #A10037) anti-Rabbit Secondary Antibody, Alexa Fluor 647 (Invitrogen #A32733)

#### FL2 Protein Upregulation Leading Edge Analysis

Scratched U2OS monolayers were incubated for decreasing intervals of time (3 hours, 1 hour, 30 minutes and 5 minutes) prior to fixation. After fixation, cells were stained for FL2 protein as described above. Cells on the wound edge (scratch periphery) were imaged using fluorescence confocal microscopy. For all scratch times, images were taken of U2OS cells with visible FL2 protein expression at the leading edge. For the untreated group, images were taken of U2OS cells in an untreated monolayer with small gaps in the monolayer to allow for background subtraction.

Fluorescence intensity at the leading edge was compared with fluorescence intensity at the cortex of cells in an uninjured monolayer. FL2 protein was quantified using FIJI ImageJ software^70^. The amount of pixels that make up 5um in an image was calculated based on the distance per pixel determined by the magnification and microscope used for imaging. This number determined the pixel amount used for the paintbrush tool in ImageJ, which was then used to outline the leading edge of the cells on the periphery of the scratch. We define the leading edge as the first 5um of the protrusions facing the scratch.

To identify the timing of FL2 protein upregulation in U2OS cells after injury, the fluorescence intensity (INTDEN) of FL2 protein within the first 5um of the cell facing the wound was determined. The corrected total cell fluorescence value was calculated (CTCF = INTDEN - [Area of selected cell * Background mean gray value]) for the intensity of FL2 protein at the leading edge of each cell. This quantity was normalized to the CTCF value of FL2 fluorescence intensity of the cortex in U2OS cells in an uninjured monolayer.

### Translation inhibition

U2OS cells were seeded at 80,000 cells per well onto glass coverslips in a 24 well plate 48 hours before treatment. Cells were treated with media containing 200ug Puromycin (Sigma P9620) or normal media as a control and incubated for 15 minutes under normal culture conditions prior to injury. A scratch was performed in the media without aspiration, PBS washes or new media addition to ensure that cells were exposed to a consistent concentration of the translation inhibitor.

The cells were incubated under normal culture conditions for one hour prior to fixation and FL2 protein staining as described below.

To determine if translation inhibition decreased FL2 protein content at the leading edge, cells with significant fluorescence intensity at the leading edge (first 5 um of cell protrusion) were counted. Fluorescence confocal images were taken of the scratch periphery with focus on areas that had cells with FL2 upregulation at the leading edge. Significant fluorescence intensity was defined as pixel intensity value at least 40% greater than that of the remaining cytoplasm of the same cell and spanned at least a fourth of the edge length. A percentage of cells with significant FL2 fluorescence intensity at the leading edge was determined for each image (number of cells with significant fluorescence intensity per image / total number of cells per image).

### Single molecule fluorescence *in situ* hybridization (smFISH)

#### U2OS smFISH

smFISH was performed according to the previously published method from Wheat, et al., 2020^42^. Cells were rinsed with PBSM (PBS with 1mM MgCl2) and fixed with 4% PFA in PBSM at room temperature for 10 minutes. Cells were washed with 50mM Glycine in PBSM for 5 minutes at room temperature. Cells were permeabilized for 10 minutes with 0.1% Triton X 100 in PBSM at room temperature and then washed with PBSM for 5 minutes thrice. Cells are then incubated at 37C for 30 minutes in a 30% pre-hybridization solution (30% formamide in 2X SSC). Hybridization of primary probes was done overnight at 37C in a 30% hybridization buffer (30% formamide, 10% dextran sulfate, 2mM VRC, 1mg/mL E. coli tRNA, 200ug/mL BSA in 2X SSC) with 100ng per coverslip of primary FL2 FISH probes and 1ul/mL Superase. The next day, cells are incubated twice for 15 minutes at 37C in 30% pre-hybridization solution prior to a five minute wash with 2XSSC and refixation with 4% PFA in PBSM for 10 minutes at room temperature. Cells are washed thrice for 5 minutes each in 2X SSC and then incubated for 30 minutes at 37C in 10% pre-hybridization solution (10% formamide in 2X SSC). Hybridization of secondary probes was done for 3 hours at 37C in 10% hybridization solution (10% formamide, 10% dextran sulfate, 2mM VRC, 1mg/mL E. coli tRNA, 200ug/mL BSA in 2X SSC) with 10ng per coverslip of fluorescently labeled secondary probe and 1ul/mL Superase. Cells are washed twice in 10% pre-hybridization solution for 15 minutes at 37C. Cells were washed thrice for 5 minutes in 2X SSC at room temperature. Coverslips were mounted using ProLong Diamond Antifade Mountant with DAPI.

#### Neuron smFISH

Protocol for smFISH in neurons was performed using the optimized smFISH protocol^42^ described above with the exceptions of the procedures indicated below.

Washes were performed with PBS-MC (PBS with 1mM MgCl2 and 0.1mM CaCl2). The fixation was performed in a solution of 4% PFA in PBS-MC.

#### smFISH-IF

FISH-IF protocol was adapted from Wu, et al., 2016, Eliscovich, et al., 2017 and the optimized smFISH protocol described above^33, 42, 44^. The procedure is the same as described above except for the differences described below.

Permeabilization was done for 15 minutes at room temperature with 0.1% Triton X 100 and 0.02% UltraPure BSA (Invitrogen #AM2616) in PBSM. Initial 10% pre-hybridization incubation was done with 10% formamide and 0.02% UltraPure BSA in 2X SSC. The 10% hybridization solution contained 1:250 GCN4 Suntag primary antibody along with the secondary probes. After hybridization, two 5 minute washes were performed at 37C with 10% pre-hybridization without BSA. The cells were incubated twice for 20 minutes with 1:800 secondary antibody (Alexa Fluor 647) at 37C.

#### Optimized smFISH Probe Design

Probes were designed using the automated design tool described in Tsanov. et al., 2016^71^. Gene specific probes are listed in Tables 1 & 2. Readout probe design and labeling were performed as in Wheat et al, Nature 2020^42^. RO6 Readout probe (GTACCGTAGGATCTGATGAA) was dual end labeled with CY3 and used for all smFISH experiments. Traditional Stellaris probes were designed on the Stellaris website using the CDS and 3’ UTR of the FIGNL2 gene (Table 3).

**Table 1.**
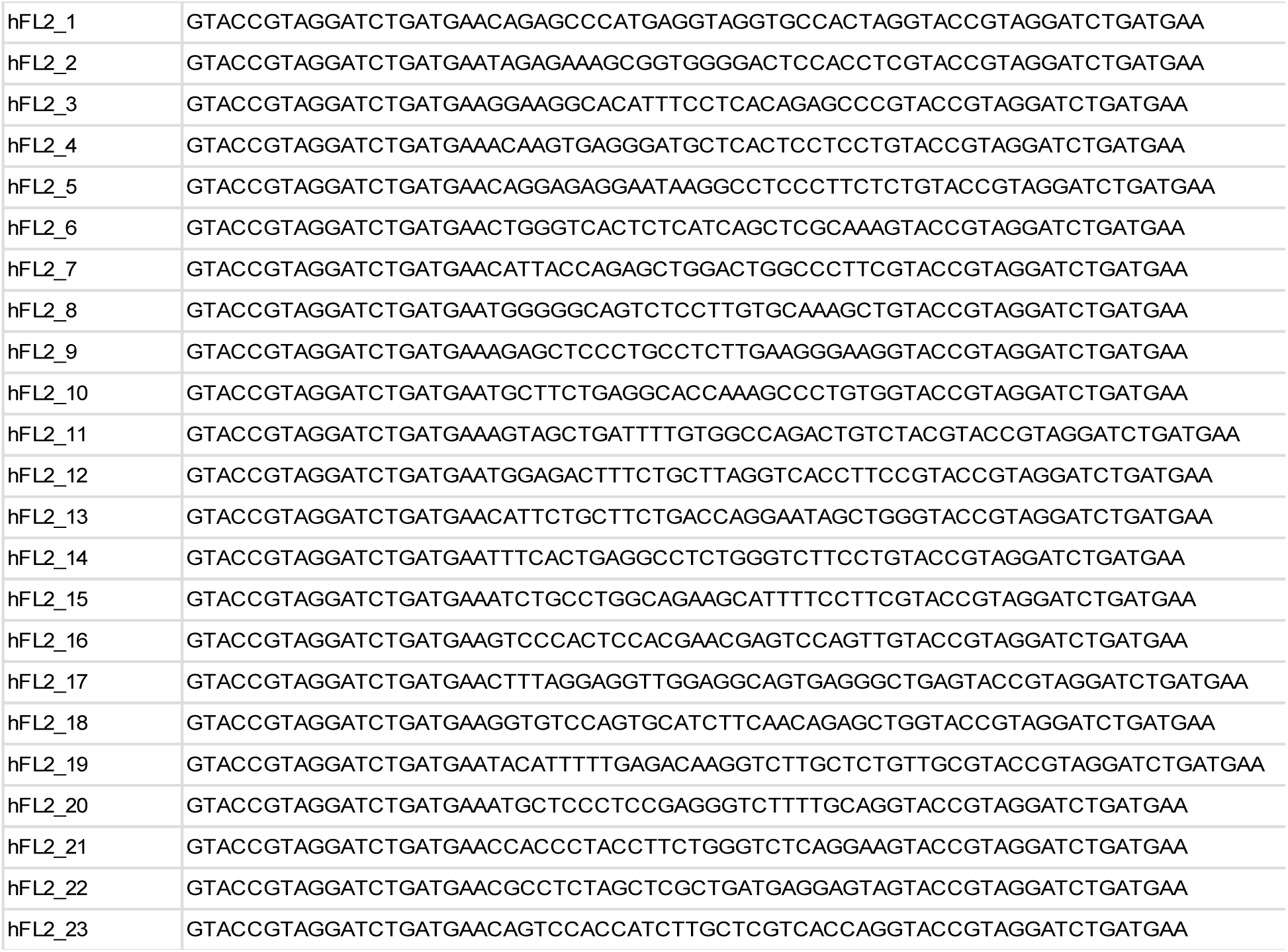
Optimized human FIGNL2 probes

**Table 2.**
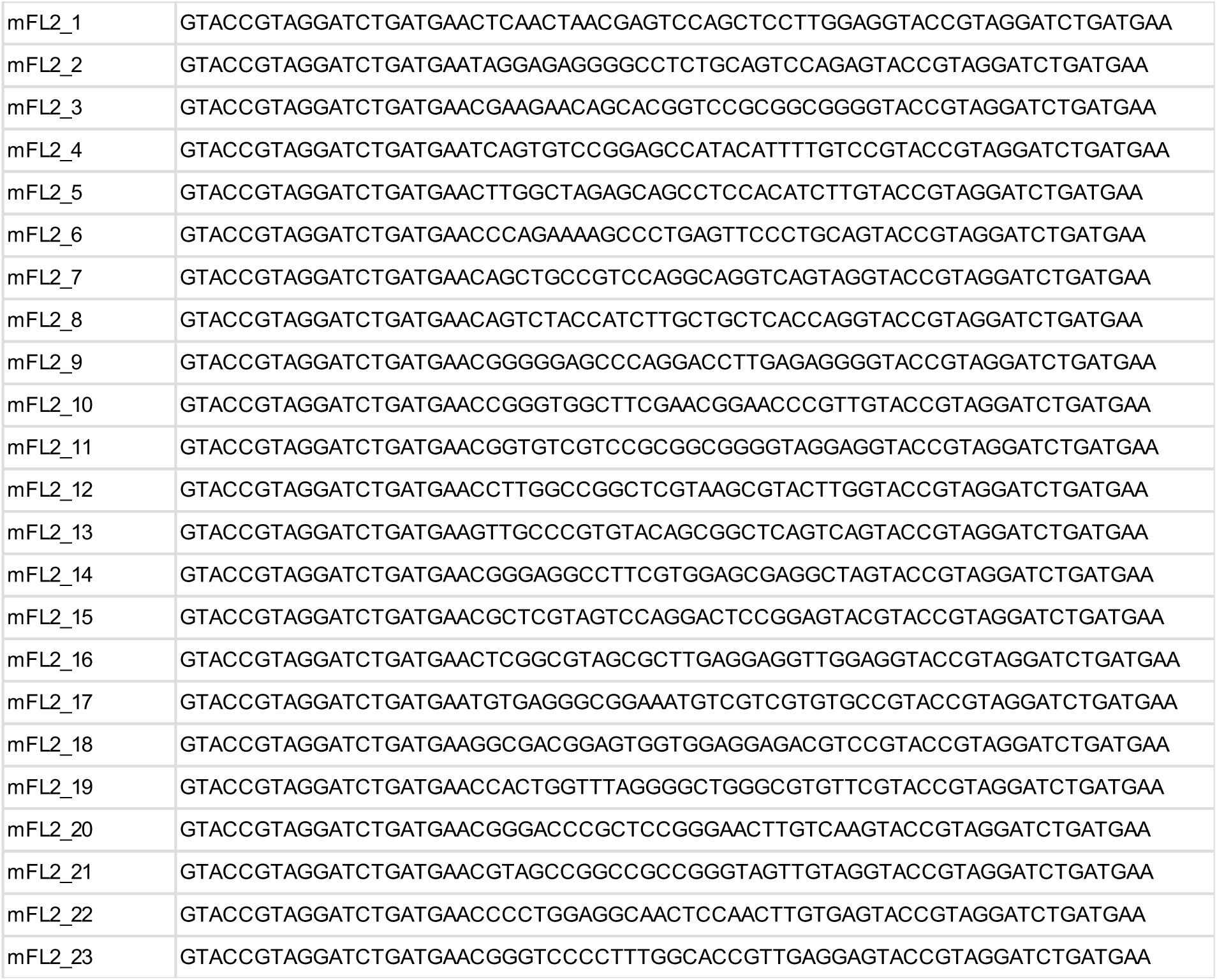
Optimized mouse FIGNL2 probes

**Table 3.**
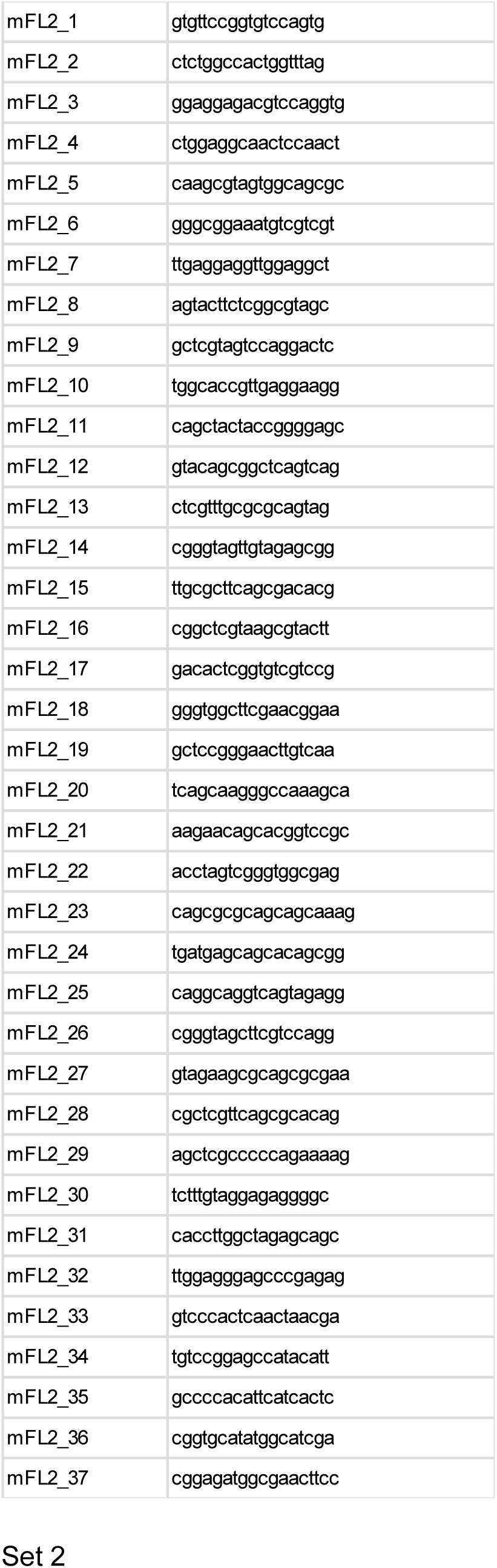
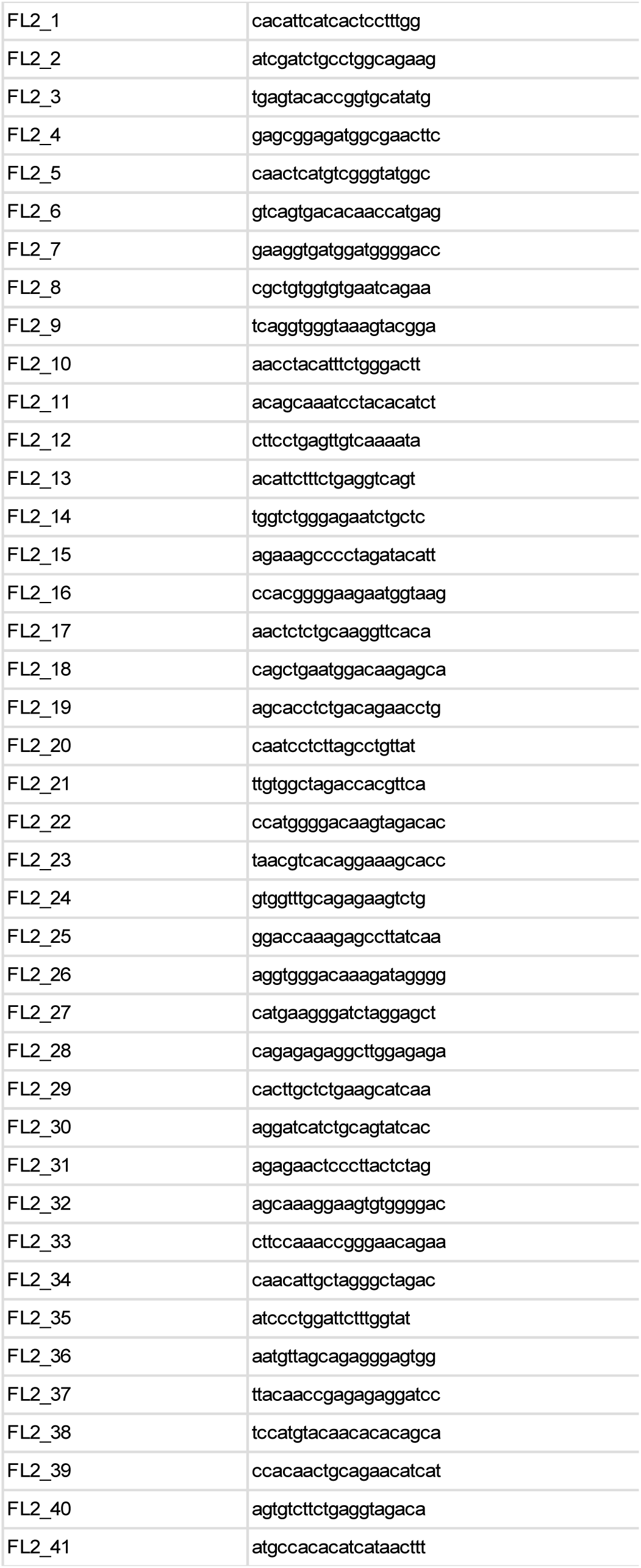
Traditional Stellaris probes Set 1

#### Leading Edge FL2 mRNA analysis

For FL2 mRNA leading edge analysis, images were taken of cells on the wound periphery of U2OS cells after one hour incubation following scratch injury. As shown previously with other localized mRNAs, not all cells have mRNA localization^3^; hence, only cells with visible FL2 mRNA localization were imaged. After imaging, mRNA molecules were identified in the images using the FISH Quant software in MATLAB^72^. Focused max projections on the z-stack images were performed for each image using the FQ_seg command. Using the max projection images, background subtraction was performed and the FISH spots were fitted to a 3D Gaussian to determine the placement of the FL2 mRNA molecules within the cell. Once the mRNA molecules were identified, the leading edge of the cells was determined in ImageJ based on our definition of leading edge (first 5um of the cell protrusion). The amount of pixels in 5um was determined for the magnification of the image and the paintbrush feature in ImageJ was used to determine how many of the identified FL2 mRNA molecules were within this compartment. The FL2 mRNA molecules that were encompassed within the leading edge outline were counted and compared to the total amount of FL2 mRNA molecules identified by FISH Quant within that image.

#### Polarization Index and Dispersion Index (PI/DI) Analysis

All image stacks were max projected and a binary mask was generated for each cell using the background autofluorescence. Then masks and corresponding max projected image stacks were run through the previously published ImageJ plugin^43^. In brief, the ImageJ plugin calculates the centroid of the cell (the mean value of x- and y-coordinates within each of the cell masks). It then subtracts the median intensity value within each cell boundary to remove background and determine an intensity weighted centroid of the RNA. The centroid of the cell and the centroid of the RNA signal are then used to calculate the RNA polarization vector. By dividing the size of the polarization vector by the radius of gyration of the cell, the polarization index is calculated. The intensity-weighted centroid of the RNA signal and the second moment are calculated to give the dispersion index.

#### Translation Site Analysis

U2OS cells were transfected with the SunTag:FL2 construct (Fig. 5A) as described above. Cells were fixed 12 hours after transfection. FISH-IF was performed on the coverslips as described above. Epifluorescence images were taken at 100X of cells with expression of the SunTag:FL2 construct. Only cells with an amount of mRNAs that was countable by eye were included in the translation site analysis (10 cells). The mRNAs and GCN4 (SunTag) proteins were identified using the FISH Quant software as described above. Using the FQ_DualColor command in FISH Quant, colocalization sites with a range of up to 600nm were identified. The paintbrush tool in ImageJ was used to identify the perinuclear area (defined as the encompassing area 5um out from the edge of the DAPI nuclear stain) and the edge area (remaining cell area outside of the perinuclear area). The number of mRNAs identified as colocalization sites were quantified in each area and divided by the total number of mRNAs in that area to obtain a percentage of translating mRNA for each region. To obtain a ratio of translating mRNA at the edge, the percent of translating mRNA at the edge was divided by the percent of translating mRNA in the perinuclear region.

### Imaging

#### Confocal Imaging

Images of fixed cell, fluorescent FL2 protein stains were taken using a spinning-disk Inverted Nikon ECLIPSE Ti-E at either 100X oil or 60X oil with numerical apertures of 1.45 and 1.4 respectively. Excitation lasers used were 405nm, 488nm and 561nm and image detection was performed via Two ORCA-FLASH 4.0 sCMOS cameras. The software used for this microscope was Nikon Elements.

#### Epifluorescence Imaging

Fixed cell images of smFISH (FL2 mRNA) and FISH-IF (FL2 mRNA and SunTag protein) were taking using an upright, wide-field Olympus BX-63 Microscope with oil immersion lenses at magnifications of either 60X (FL2 mRNA images) or 100X (FISH-IF images) and numerical apertures of 1.35 and 1.4 respectively. An ORCA-R2 digital CCD camera was used with DAPI, Cy3 and Cy5 filters from Semrock. The software used for image acquisition and stage control was Metamorph (Molecular Devices).

### Cloning and PCRs

#### Plasmids and cloning

For cloning of the SunTag-FL2 construct, we used the SINAPs plasmid from the Singer lab (Addgene #84561)^44^ and a tdTomato-FL2 (containing the 3’UTR of FL2) plasmid previously cloned in-house using the tdTomato-C1 vector (Addgene #54653), a human FL2 clone in pANT7_cGST (DNASU, Arizona State University, Tempe, AZ, clone: HsCD00403041) and a human FL2 3’UTR construct (Switchgear Genomics #S811553). The Ubc promoter, flag tag and SunTag reporter were cloned from the SINAPs construct to replace the CMV promoter and tdTomato reporter of the tdTomato-FL2 plasmid using NEBuilder HiFi DNA Assembly technology (NEB).

The plasmids used for IMP RBP overexpression, GFP-ZBP1^33^, GFP-IGF2BP2 and GFP-IGF2BP2 KH3 mutant^39^, were from the Singer laboratory. The Lin28 plasmids were purchased from Addgene; Lin28A (#51371), Lin28B (#51373) and Lin28A-mCCHC (#51372). The miRNA let-7 inhibitor sequence (3’-CUCCAUCAUCCAACAU-5’) was previously validated for broad let-7 family inhibition (Frost & Olson, 2011). The custom miRCURY LNA miRNA inhibitor was purchased from Qiagen (Cat num 339146).

#### RT-PCR

Since identification of FL2 levels within human cell lines via qPCR has been so far unsuccessful, quantitative mRNA analysis was carried out in the neuroblastoma Neuro2A cell line. Cells were transfected with the indicated plasmid or inhibitor as described above and incubated for 48 hours prior to collection. Collection and cDNA synthesis were done using the SuperScript IV Cell Direct cDNA Synthesis Kit (Thermo Fisher #11750150) and carried out according to the kit protocol. *Power* SYBR Green PCR Master Mix was used for comparative Ct qPCR run on the ViiA7 Real-Time PCR System. Predesigned PrimeTime qPCR Primers were purchased from IDT. FL2 primers, FignL2 Assay ID: Mm.PT.58.21940655.g; GAPDH control primers, GAPDH Assay ID: Mm.PT.39a.1. Results were analyzed using the 2^(ΔΔCt) method. The caveat is that these were transiently transfect cells, so the amount of cells transfected from one experiment to the next is undetermined. Averaged results from 4 independent experiments.

## LEAD CONTACT

Further information and requests for resources and reagents should be directed to the lead contact, David Sharp.

## MATERIALS AVAILABILITY

All FISH probes designed and used in this study are listed in the supplementary tables. Constructs generated for this study are available upon request to the lead contact.

## DATA AND CODE AVAILABILITY

All raw imaging data generated for this study is available upon request to the lead contact.

---

**Supplemental Figure 1.**
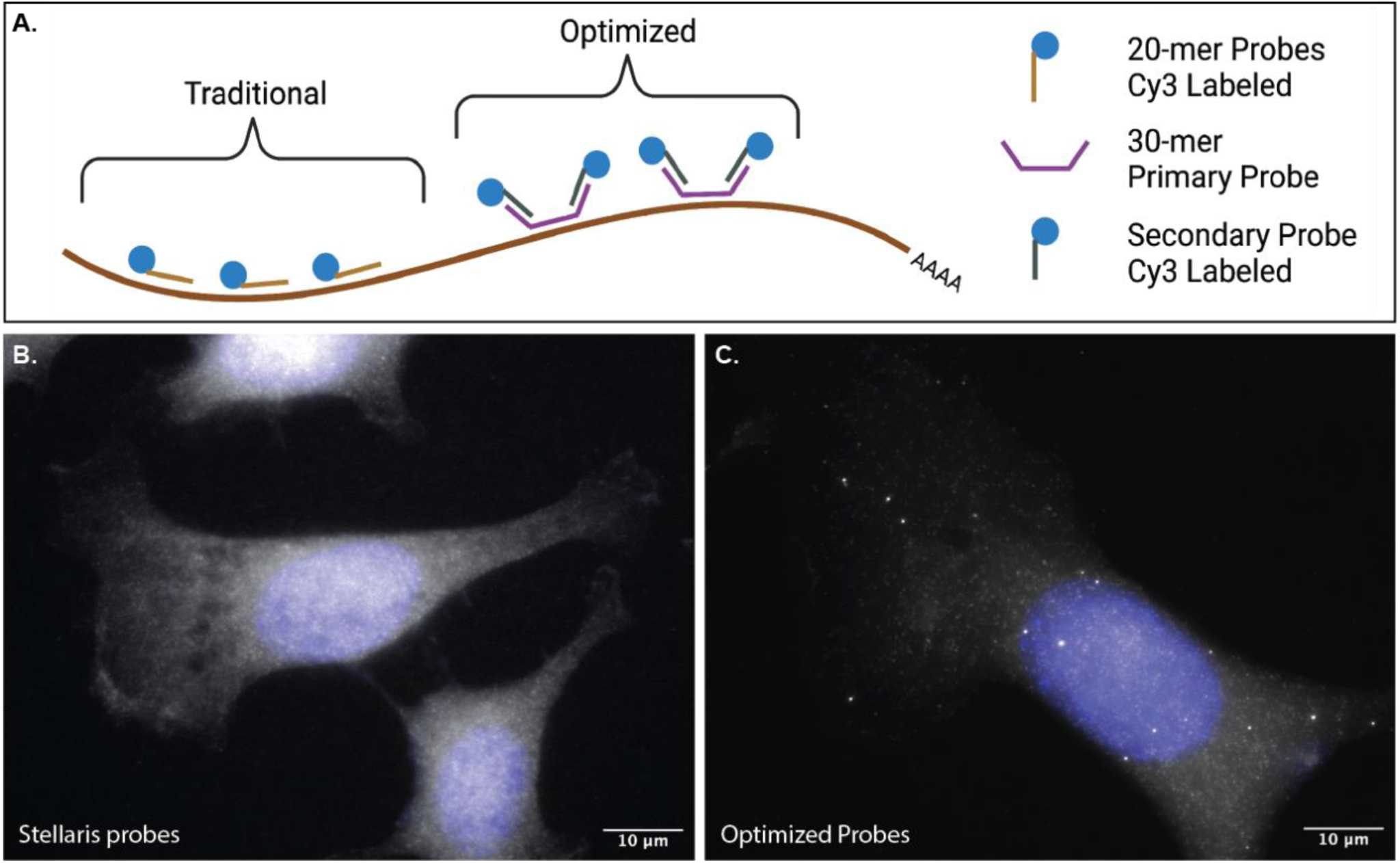
Optimization of smFISH for visualization of FL2 mRNA. A) Schematic of the difference between the use of traditional Stellaris smFISH probes and optimized smFISH probes. The Stellaris probes use fluorescently labeled 18-22mer probes that hybridize to the mRNA sequence. The optimized probes use a primary set of 30mer probes that hybridize to the mRNA sequence and a secondary set of fluorescently labeled probes that hybridize to the 20mer overhangs of the primary set. B) FL2 mRNA smFISH in MEFs using Stellaris probes. C) FL2 mRNA smFISH using optimized probes.

**Supplemental Figure 2.**
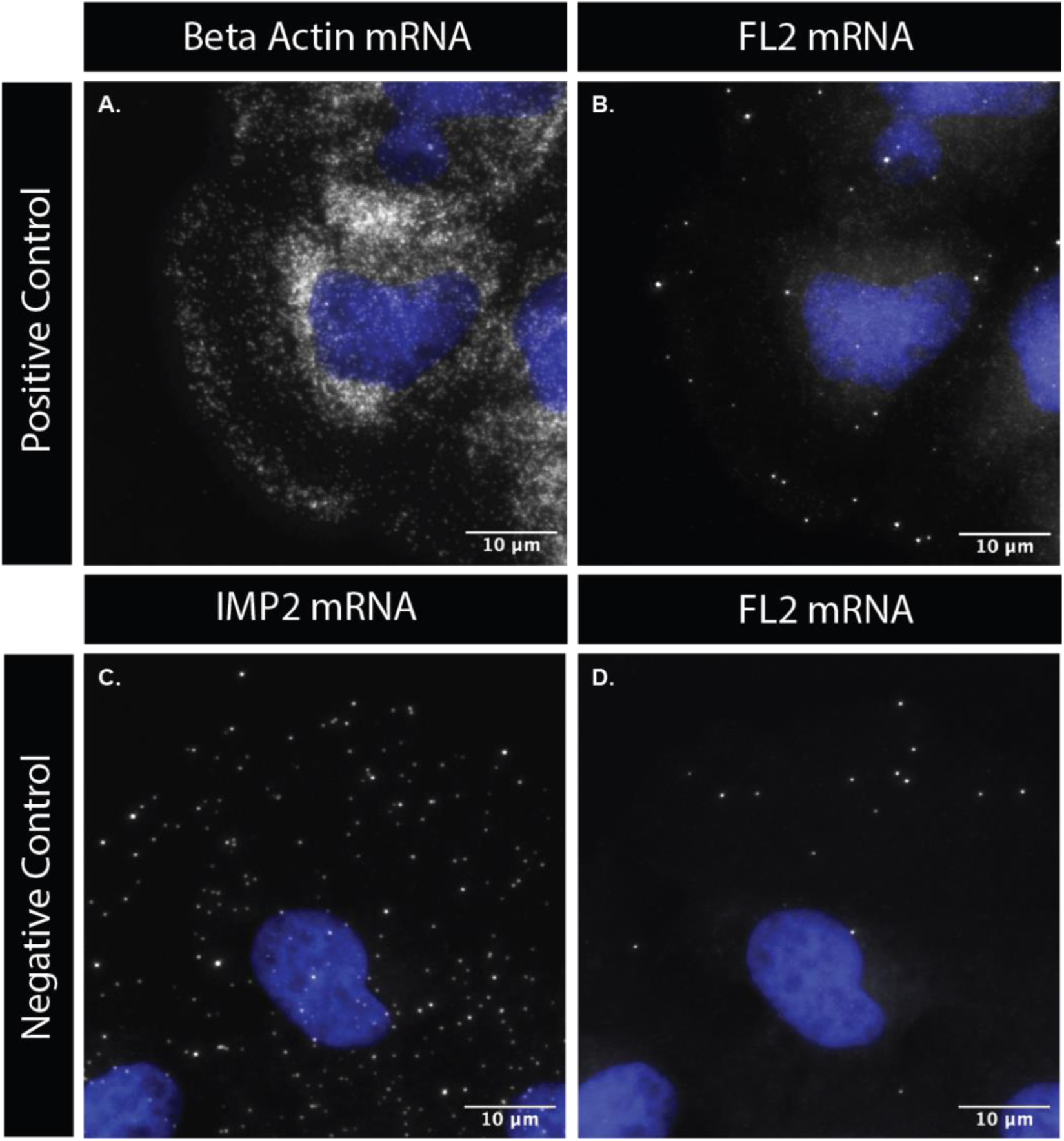
Comparative smFISH images of control mRNA and FL2 mRNA used for the polarization and dispersion index assay. A) β-actin mRNA used as the positive control. B) FL2 mRNA in the same cell used in panel A. C) IMP2 mRNA used as a negative control. D) FL2 mRNA in the same cell used in panel C.

**Supplemental Figure 3.**
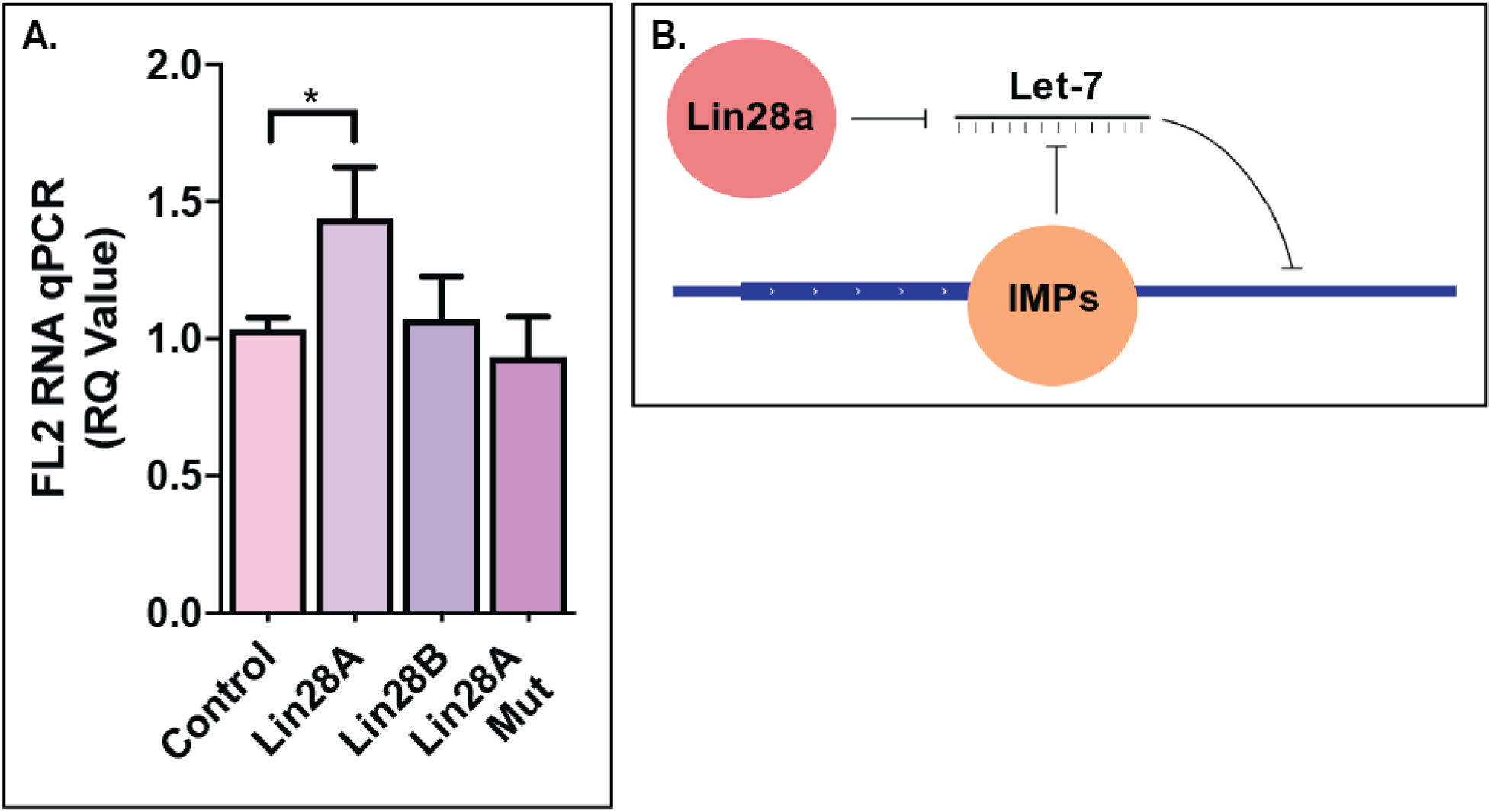
LIN28 indirectly affects FL2 mRNA expression by negatively regulating let-7 miRNA expression. A) FL2 mRNA quantity in N2A cells with 48 hours of LIN28 overexpression. Lin28A Mut is a mutant that cannot negatively regulate let-7 miRNA. B) Schematic of the regulating molecules involved in FL2 mRNA expression.

**Supplemental Figure 4.**
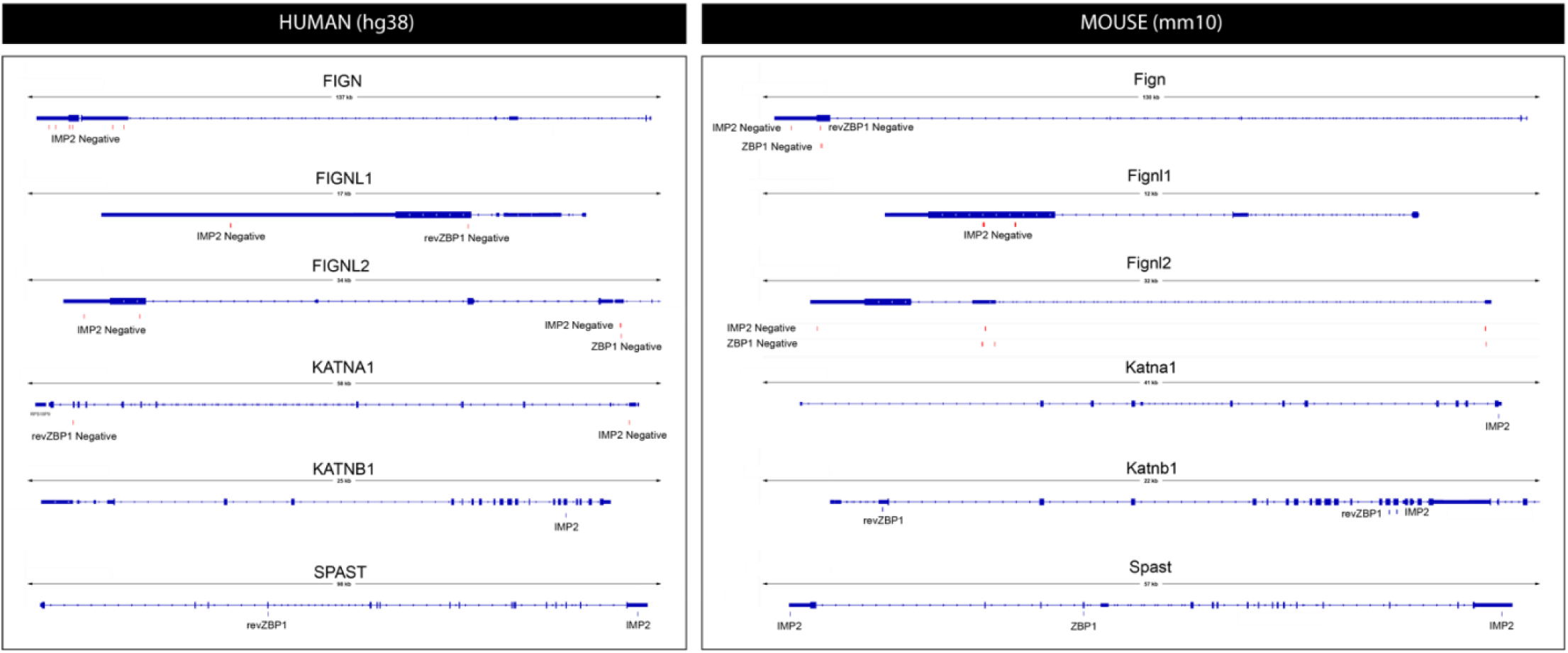
The IMP RBPs also have the capacity for binding to other MSEs. IMP1 (ZBP1) and IMP2 RBP binding motifs within the human (left) and mouse (right) gene sequences of the MSEs, including fidgetin (FIGN, Fign), fidgetin-like 1 (FIGNL1, Fignl1), fidgetin-like 2 (FIGNL2, Fignl2), katanin p60 (KATNA1, Katna1), katanin p80 (KATNB1, Katnb1) and sptastin (SPAST, Spast). IMP1/ZBP1 motifs from (Patel et al, Genes and Development 2013) and IMP2 motifs from (Biswas et al, Nat Com 2019). Positive and negative correspond to DNA strand on which the motif search was performed (all strands correspond to the gene of interest). Motif search limited to exons (thick lines) only, motifs found in introns (thin lines) were excluded as they were unlikely to be present in the mature mRNA.

## Notes

### Competing Interest Statement

The authors have declared no competing interest.

